# Differences in fMRI-based functional connectivity during abstinence or interventions between heroin-dependent individuals and healthy controls

**DOI:** 10.1101/2024.06.11.598479

**Authors:** Danielle L. Kurtin, Anusha M. Prabhu, Alissa Groen, Matthew J. Amer, Anne Lingford-Hughes, Louise M. Paterson

## Abstract

The substantial personal, societal, and economic impacts of opioid addiction drive research investigating how opioid addiction affects the brain, and whether therapies attenuate addiction-related metrics of brain function. One useful approach to characterise the effects of opioid addiction on the brain is functional connectivity (FC). FC assesses the pairwise relationship of brain region function over time. This work is a systematic narrative review of studies investigating the effect of abstinence or interventions on FC in people who are dependent on heroin (HD) and healthy controls (HC). We found that HD typically showed weaker FC between three functional networks: the Executive Control Network, Default Mode Network, and the Salience Network. Abstinence and Transcranial Magnetic Stimulation (TMS) both attenuated differences in FC between HD and HC, often by increasing FC in HD. We critically assessed the clinical relevance of these results and the impact of study methods on the robustness of study results. We concluded with practical suggestions to improve the translational potential of neuromodulatory interventions (e.g., noninvasive brain stimulation) targeting the neural correlates of opioid addiction.

**Highlights:**

- Functional connectivity (FC) informs how neural resources are organised.
- FC is weaker in people with heroin dependence (HD) vs healthy controls (HC)
- Weaker FC is observed among resting, reward, cognitive, and attentional networks.
- Abstinence and brain stimulation strengthen FC and reduce craving/relapse in HD.
- FC may be a biomarker for developing therapies for HD.

## 1. Introduction

Illicit drugs were estimated to cost the UK £20 billion in 2020 (Black, 2021) and $1.02 trillion in the US in 2017 (Florence et al., 2021). The current cost of addiction is likely growing, given that drug addiction showed the first and second-highest increase in percent change of disease burden between 1990-2019 for women and men, respectively (McKee et al., 2021). Out of all drugs of abuse, opioids were the largest contributor to the record high in drug-related mortality in 2018 in the UK (Black, 2021). In the US, nearly 135 people died per day in 2019 from opioid-related deaths (Center for Disease Control and Prevention, 2022). The substantial health, social, and economic impacts of opioid addiction necessitate research of the neural correlates of opioid addiction, and whether they are attenuated by therapeutic interventions.

As with all addictions, opioid addiction is characterised by chronic cycles of relapse and remittance, which relate to dysfunctional neural circuitry underpinning impulse control, decision-making, emotional regulation, stress response, and reward processing (Hayes et al., 2020; Koob, 2009; Kozak et al., 2019; Kwako et al., 2016; Moeller & Paulus, 2018; Uhl et al., 2019; Wilcox et al., 2019). Effective therapies that result in maintained abstinence have been shown to restore or attenuate these altered neural circuits (Koob, 2006; Koob & Volkow, 2010; Lueptow et al., 2020). Thus, characterising how the neural correlates of opioid addiction are affected by abstinence or therapeutic interventions will help improve the development of therapeutic interventions.

### 1.1 Identifying the neural correlates of opioid addiction with neuroimaging

One way to characterise the neural correlates of opioid addiction is through functional magnetic resonance imaging (fMRI), by utilising the Blood Oxygenation Level Dependent (BOLD) signal as a proxy measure of neuronal activity (Detre & Wang, 2002). FMRI acquired during resting state captures spontaneous, intrinsic brain activity, whereas task-fMRI is used to probe the neural correlates of specific processes. For example, cue-reactivity tasks show blocks of drug-related images, followed by blocks of neutral visual stimuli. People with opioid addiction tend to show higher BOLD activity in reward regions as compared with healthy controls (HC) in response to drug-related stimuli (Hayes et al., 2020).

Another way to use fMRI to assess the neural correlates of opioid addiction is to use Functional Connectivity (FC). FC analyses evaluate the relationship between two or more brain regions, often via the temporal correlation between spatially separated brain regions (Biswal et al., 1995; Fingelkurts et al., 2005). There are many ways to compute FC, each with trade-offs and considerations. In Box 1 we consider how the method by which researchers define brain regions impacts FC analysis. In Box 2, we provide an overview of common FC methods.

#### Box 1. Defining brain regions.

The first step of most FC analyses is to identify the regions from which timeseries of BOLD activity are extracted and compared. Dividing the cortex into discrete regions is known as parcellation, which can be broadly split into anatomical or functional approaches. Anatomical parcellation uses atlases that define brain regions based on morphological, structural, or cytoarchitectural characteristics (Amunts & Zilles, 2015). Functional parcellation defines brain regions by processes in which it is typically associated (Eickhoff et al., 2018). For example, one of the earliest fMRI studies presented participants with a checkerboard visual stimulus, and in response, the posterior occipital cortex exhibited an increased magnitude of the BOLD signal, associating its function with vision (Kwong et al., 1992). It is best practice to use functional parcellation (especially for cortical regions) for FC analyses, as regions defined anatomically may have functional variation and thus obscure the ability to assess the role of the region in the function under evaluation (Bijsterbosch et al., 2020). Within functional atlases, brain regions can be defined either by discrete boundaries, referred to as hard parcellations, or with non-distinct boundaries, known as soft parcellations. One consideration for atlas-based parcellations is whether to parcellate data in standard vs subject space. The former is more common, as registering subject’s functional data to standard space is often included in preprocessing, and subsequent parcellation using an atlas in standard space is straightforward. However, distortions are inevitable when registering subject data to standard space, resulting in heterogeneous changes in regional volume, and potential misalignment of spatial boundaries (Allen et al., 2008). This negatively impacts FC analysis, since average regional timeseries will be influenced by voxels inappropriately grouped within a region (Bijsterbosch et al., 2019; S. M. Smith et al., 2011). By conducting analyses in subject space, or by comparing FC results from standard space to those in subject space, researchers can be more confident their results are not driven by artefacts associated with spatial registration. A more advanced method of subject-level parcellation is to conduct individual functional parcellation. This approach uses participant’s neuroimaging data to inform personalised boundaries between regions (Kong et al., 2019; Laumann et al., 2015; D. Wang et al., 2015). Atlas-based parcellations are not the only way to define regions from which FC is quantified; studies may also define regions based on their engagement during conditions of interest (e.g., blocks of drug-related stimuli in a cue reactivity task). For example, a study may identify a region related to dysfunctional reward processing in people with opioid addiction based on a meta-analysis’s identification of voxels that show reliable group differences during a cue reactivity task. Another approach is to employ Independent Component Analysis (ICA), which identifies voxels or regions that are reliably engaged in a particular condition (Duff et al., 2012).

#### Box 2. How is functional connectivity defined?

There is a large variety of methods to compute FC, which can be broadly characterised using two axes - directed vs undirected, and model-based vs model-free (Farahani et al., 2019; K. Li et al., 2009; Rogers et al., 2007). Both directed and undirected FC approaches assess whether two timeseries are related; however, directed measures also capture whether the relationship is driven by one region’s influence over another. A common example of an undirected FC approach is correlation between two timeseries, where a correlation coefficient indicates the strength of interregional relationships, but not whether one region is influencing another. Common methods for directed connectivity, also known as effective connectivity, include Dynamic Causal Modelling (Friston et al., 2003), Granger Causality (Goebel et al., 2003), and Transfer Entropy (Schreiber, 2000; Sharaev et al., 2016). Model-based methods are the more common form of FC analysis. They work by extracting the timeseries of BOLD activity from a brain region and computing the relationship between other regional timeseries. A common model-based FC approach is to compute the correlation or coherence between pairs of timeseries (Biswal et al., 1995; Greicius et al., 2003; Wig et al., 2011). Psychophysiological interactions (PPI) are another model-based approach that works by modelling the influence of task conditions on pairwise connectivity (Cisler et al., 2014; Friston et al., 1997; O’Reilly et al., 2012). Model-free FC approaches are data-driven, and do not consider regional specificity or task conditions - instead, model-based methods typically cluster or reduce the dimensionality of the data. Clustering-based FC is less common than dimensionality reduction approaches, and for more information, see the following references (Cordes et al., 2002; Golay et al., 1998; Seghier, 2018). Dimensionality reduction approaches include principal component analysis (PCA) and independent component analysis (ICA), which identify components of spatial or temporal connectivity (Calhoun et al., 2009). While this section provides a snapshot of the breadth of FC approaches, there is little consensus as to which approaches are superior. Direct evaluation of both model-based vs model-free (Baumgartner et al., 1998; Worsley et al., 2005) and directed vs undirected (Conti et al., 2019) approaches showed comparable performance. Since each FC method has trade-offs and limitations, it is sensible to choose an approach based on the aims of each study (K. Li et al., 2009). For example, model-based methods facilitate targeted exploration of how regions interact with other regions, whereas a model-free approach is unconstrained by assumptions underpinning parcellation; undirected FC methods can tractably characterise how all pairs of regions coordinate during a condition, whereas directed FC methods are more useful in establishing a hierarchy of function among a group of regions (Ray et al., 2021; Rolls et al., 2022; Vaudano et al., 2009).

FC analysis is especially useful for understanding how brain regions work together to subserve complex, distributed processes impaired in addiction. Here we first describe the role of functional networks, and their relationship to studies investigating differences between people with opioid addiction and HC.

### 1.2 The role of functional networks

Groups of regions that reliably coactivate or connect in a particular condition are deemed networks. For example, there are networks associated with different sensorimotor processes, such as the visual, somatosensory, auditory, and motor networks. Other notable networks support attention (e.g., dorsal attention network and Salience Network (SN)), reward processing (e.g., limbic and ventromedial networks), cognition (e.g., executive control network (ECN)), and resting state (Default Mode Network (DMN)) (Schaefer et al., 2018; Yeo et al., 2011). Investigating network topology (i.e., how/what regions are included in a network) and dynamics (i.e., prevalence and coordination among networks across time) informs how different groups utilise process-specific neural resources. Opiate addiction has been shown to affect FC in reward processing networks, the ECN, DMN, and SN (Chen et al., 2022; Dunlop et al., 2017; Jin et al., 2022; Q. Li et al., 2018). For example, research has shown people with opioid addiction exhibit altered FC between networks involved with stress/emotion regulation and cognition as compared with HC (Belin et al., 2013; Dugré et al., 2023; Everitt & Robbins, 2016; Fareed et al., 2017).

### 1.3 Aims and objectives

The first aim of this systematic narrative review is to characterise the different effects of abstinence or interventions on FC between HC and people with opioid addiction. Our second aim is to evaluate whether the effects of abstinence or an intervention on FC is associated with clinical/behavioural outcomes. While there are recent reviews on differences in FC between HC and people with drug addiction (Koob, 2021; Wilcox et al., 2019), our work is notable for its focus on opioid addiction. Among reviews on the neural correlates of opioid addiction, our focus on characterising the effect of abstinence or an intervention on FC in people addicted to illicit opioids vs HC is distinct. This is because reviews on the neural correlates of opioid addiction tend to include both microscale (e.g., receptor-level changes) and macroscale measures (e.g., FC) of brain function or focus on intrinsic differences between people with opioid addiction and HC (Herlinger & Lingford-Hughes, 2022; Moningka et al., 2019). Our focus on changes in FC in people with opioid addiction as a result of abstinence or an intervention increases the translational relevance of this review in three ways. First, developing an understanding of differences in FC can inform the delivery of targeted neuromodulatory, therapeutic interventions, such as noninvasive brain stimulation. Second, identifying an association between abstinence- or intervention-related changes in clinically relevant effects (e.g., decreased craving or risk of relapse) and FC provides potential biomarkers for developing therapeutic/neuromodulatory interventions. Finally, our evaluation the impact of study methods on the reliability and generalisability of results enables improved approaches of future research.

## 2. Methods

This study was preregistered on PROSPERO (CRD42023418691). A literature search was carried out in April 2023 across the following 6 electronic bibliographic databases: MEDLINE, EMBASE, American Psychological Association (APA) PsycINFO, PsycArticlesFullText, Scopus and PubMed.

Search terms relating to illicit opioid dependence, FC, and fMRI were employed as provided in Appendix 1 (Fig 1). After removal of duplicates across databases, 438 papers remained. After removing articles classified as non-English language articles, narrative reviews, systematic reviews, conference abstracts/papers, dissertations, or qualitative studies (e.g., case studies, opinion pieces, letters, comments, editorials), 266 articles remained. These articles’, title, abstract, and text were screened more closely to ascertain whether they met inclusion criteria by 3 reviewers, with 2 independent reviewers reviewing each article against the pre-defined inclusion and exclusion criteria (Table 1). This concluded in initially including 13 articles, with 33 articles that reviewers disagreed on. Disagreements about screening were resolved by discussion between all the 3 reviewers, which resulted in 9 out of the 33 articles moving to full text screening. Full text screening was performed on the 22 articles agreed upon by all 3 reviewers. Each article was reviewed by two independent reviewers using a standardised data extraction document on Microsoft Excel with predefined columns following the PRISMA protocol guidelines (Page et al., 2021). The results of data extraction were checked for consensus across the two reviewers for each article and disagreements were settled through discussion. Out of the 22 articles that underwent full text screening, 6 were determined to not meet inclusion criteria. For example, two studies included rsfMRI in people with opioid addiction and HC, but could not directly assess the effect of a task-based intervention on FC because tasks were performed outside the scanner (Q. Li et al., 2013; Ma et al., 2015).

**Figure 1:**
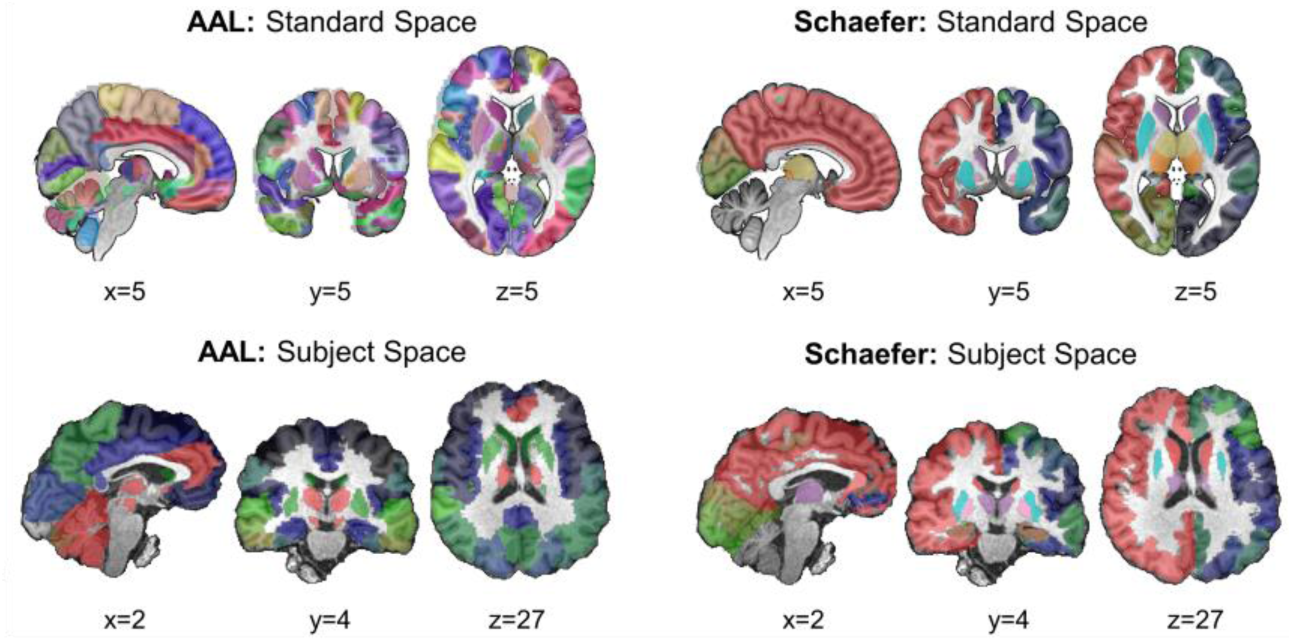
Example of (A) anatomical and (B) functional atlases. (A) The Automated Anatomical Labelling (AAL) atlas is overlaid on (Ai) a standard MNI brain and (Aii) after transformation to subject space. The Schaefer atlas in (Bi) standard MNI space and (Bii) subject space is an example of a functionally derived parcellation (Schaefer et al., 2018). In all panels, colours demarcate different brain regions. Sagittal, coronal, and transverse slices are provided by the x, y, and z coordinates, respectively.

**Table 1:**
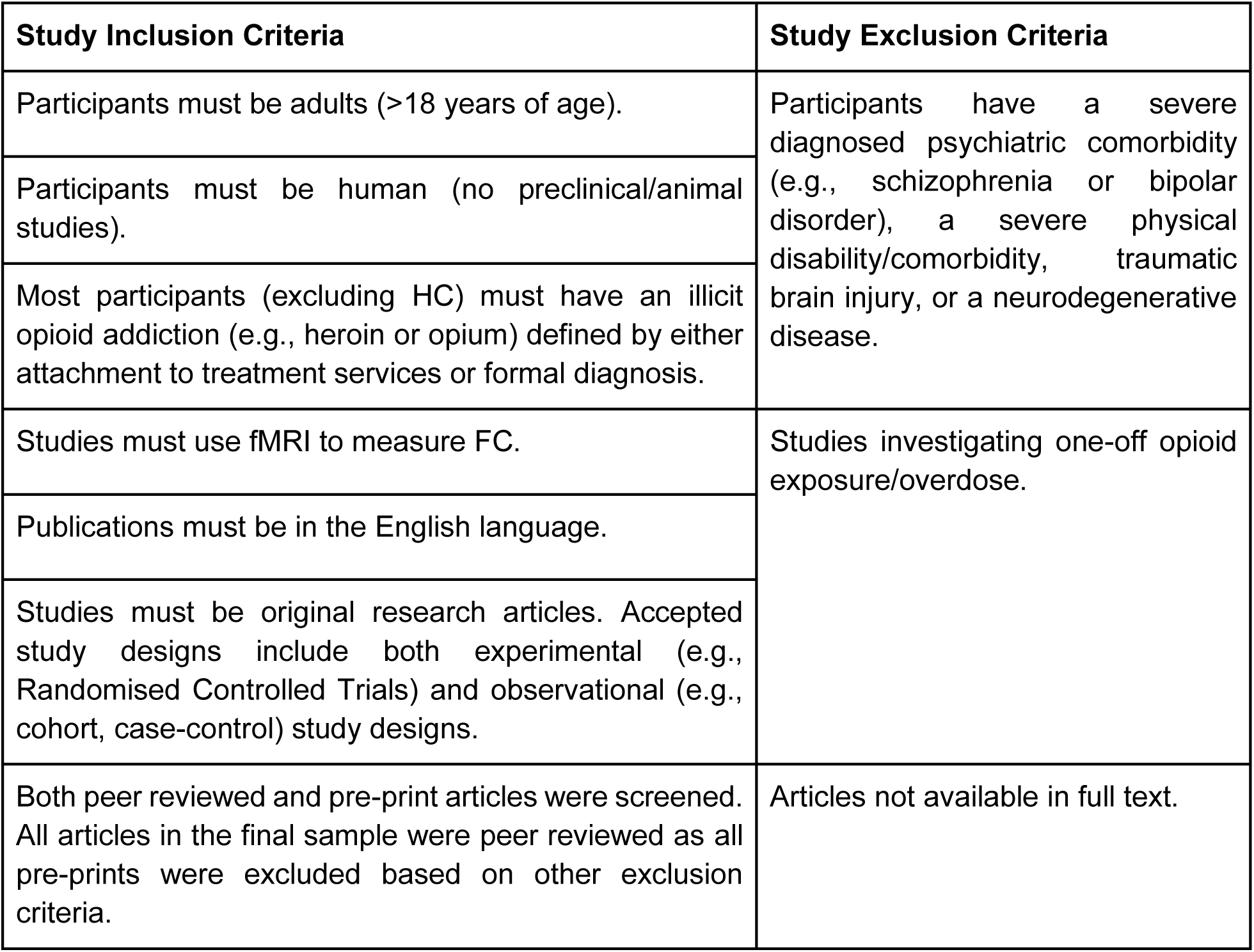
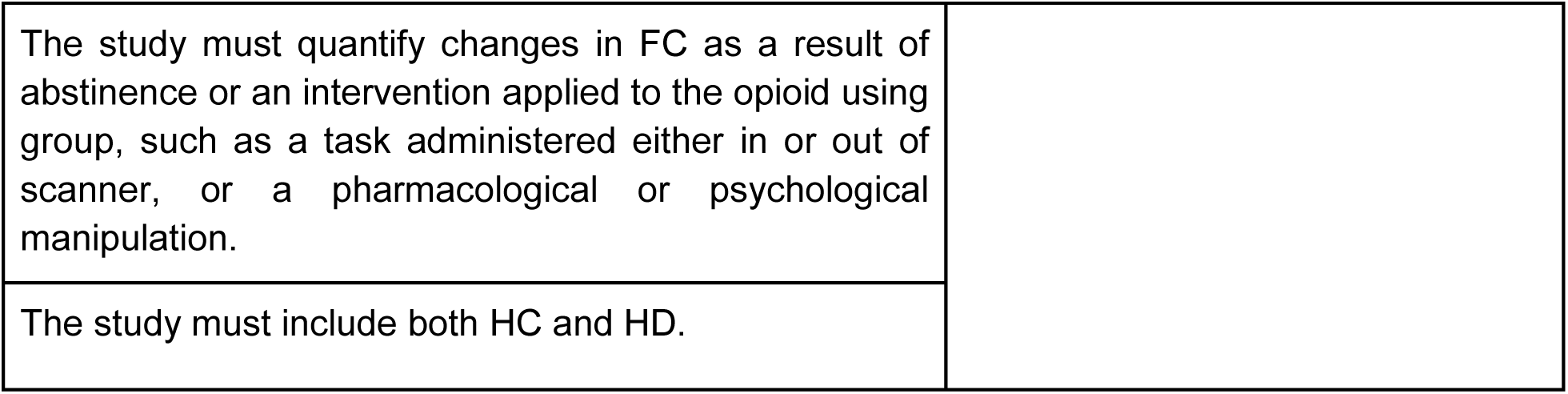
Study inclusion and exclusion criteria.

Thus, 16 articles were selected as the final dataset for qualitative, narrative synthesis of results in line with Synthesis Without Metaanalysis (SWiM) guidelines (Campbell et al., 2020) (Fig 1). All studies included individuals who were dependent on heroin; thus, groups with opioid addiction will be referred to as heroin-dependent (HD) throughout this work.

## 3. Results

### 3.1 Demographics and population characteristics

The majority of studies reviewed in this work were conducted in China (Studies 1-13, Table 2), with the remaining 3 conducted in Switzerland (Studies 14-16). The average participant age in each study ranged from 32-49 years old. Eight studies included only male participants, and all 16 studies reported no significant difference in biological sex between HD and HC groups. Study designs included 7 comparative cohort studies between HC and HD, 2 randomised controlled trials, 1 non-randomised interventional trials and 6 cross-sectional studies. Opioid use disorder or dependence was classified primarily by DSM-IV or DSM-5 criteria. One study used ICD-10 criteria to define opioid addiction, and another used a prescription of Methadone-Maintained Therapy (MMT) (Study 16). Definitions of abstinence varied across studies and a wide range of defined abstinence length was represented. Eleven of 16 studies included individuals receiving opioid substitution therapy (OST), primarily methadone.

**Table 2:**
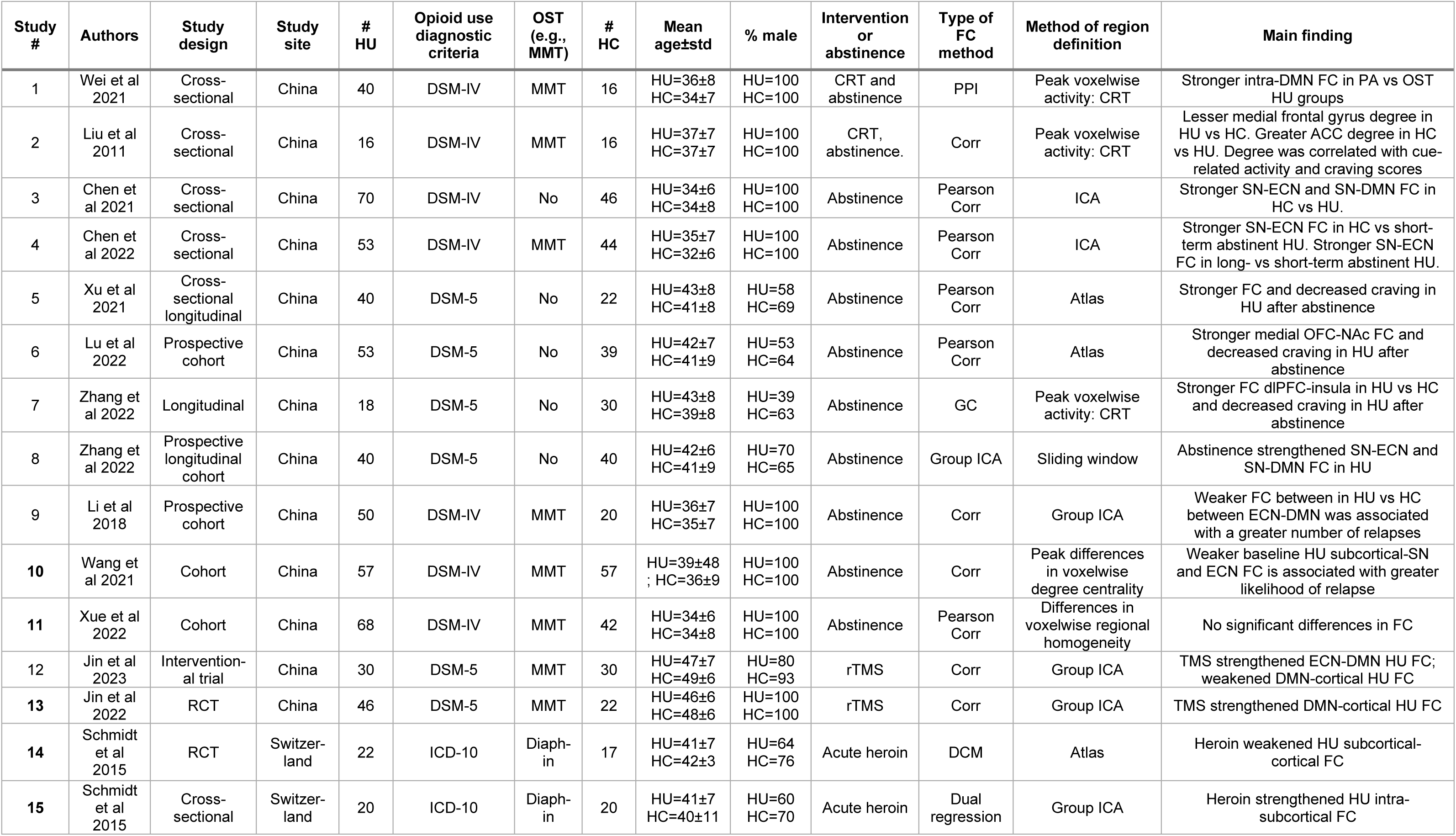
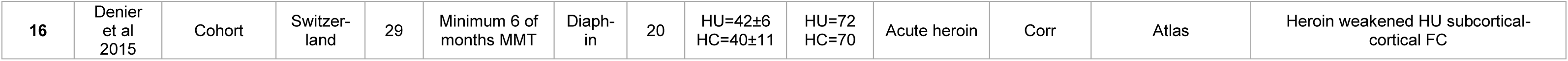
Study information. Abbreviations: RCT=Randomised Control Trial; OST=Opioid Substitution Therapy; MMT=Methadone Maintained Therapy; CRT=Cue Reactivity Task; FC=Functional Connectivity; rTMS=randomised Transcranial Magnetic Stimulation; PPI=psychophysiological interaction; Corr=Correlation; ICA=Independent component analysis; GC=Granger Causality; DCM=Dynamic Causal Modelling; PA=Protracted Abstinence; SN=Salience Network; ECN=Executive Control Network; DMN=Default Mode Network.

Most studies investigated the effects of abstinence on FC (Studies 1-11, Table 2). The remaining 5 studies utilised two other interventions; Transcranial Magnetic Stimulation (TMS, a form of noninvasive brain stimulation (studies 12-13) and acute administration of heroin (Studies 14-16).

### 3.2 Comparisons between HC and HD population changes in functional connectivity as a result of cue-reactivity tasks or abstinence

All studies that investigated the effects of either a cue-reactivity task or abstinence on FC were conducted in China (Studies 1-11, Table 2). Most collected resting state data (Studies 3-11). Of the two cue-reactivity studies, one (Wei et al., 2021) (Study 1), showed that drug-related cues recruited stronger FC between two DMN regions, the Posterior Cingulate Cortex (PCC) - medial Prefrontal Cortex (mPFC), in HD with protracted abstinence (i.e., non-OST-maintained abstinence for 6 months) as compared with people with OST-maintained abstinence (Wei et al., 2021). Given there were no significant differences in FC between HD to HC, it is unclear whether OST- or non-OST-maintained abstinence restored FC in the direction of HC, and whether that related to decreased craving.

The other study collected data during both resting state and while participants performed a cue-reactivity paradigm (J. Liu et al., 2011) (Study 2, Table 2). During resting state, the authors reported that the medial frontal gyrus had greater degree (i.e., number of connections to other regions) in HU as compared with HC. Conversely, the Anterior Cingulate Cortex (ACC) had a lower degree in HU as compared with HC. Relationships between rsfMRI degree and activity during the cue reactivity task were observed. The weakened FC in HD from the medial frontal gyrus was negatively correlated with both heroin cue-related activity in medial frontal gyrus, and craving scores. Conversely, the stronger FC in HD in the ACC was positively correlated with both cue-related activity in the ACC and craving scores (J. Liu et al., 2011).

The remaining 9 resting state studies investigated relationships between abstinence and changes in FC (Studies 3-11, Table 2). All studies included at least one HC scan for comparison to baseline and post-abstinence scans in HD. All studies included two scanning sessions for HD: one at baseline, and a second after the duration of abstinence. The duration of abstinence ranged from short (minimum = 3 days) to long term (maximum = 26 months). Moreover, the definition of abstinence varied across studies, as 5 out of 9 abstinence studies considered HD abstinent while on Opioid Substitution Therapy (OST). Despite these differences, studies consistently found HC had the strongest FC between nodes in the SN and ECN, followed by long-term abstinent HD, then short-term abstinent HD (Fig 2). This pattern of stronger FC between SN and ECN nodes as a function of abstinence occurred in 61% (74 out of 122 edges (i.e., connectivity between a pair of regions)) of the significant changes in FC associated with abstinence. Stronger ECN-SN FC as a function of abstinence was consistently observed across studies utilising HD maintaining abstinence with and without OST.

**Figure 2:**
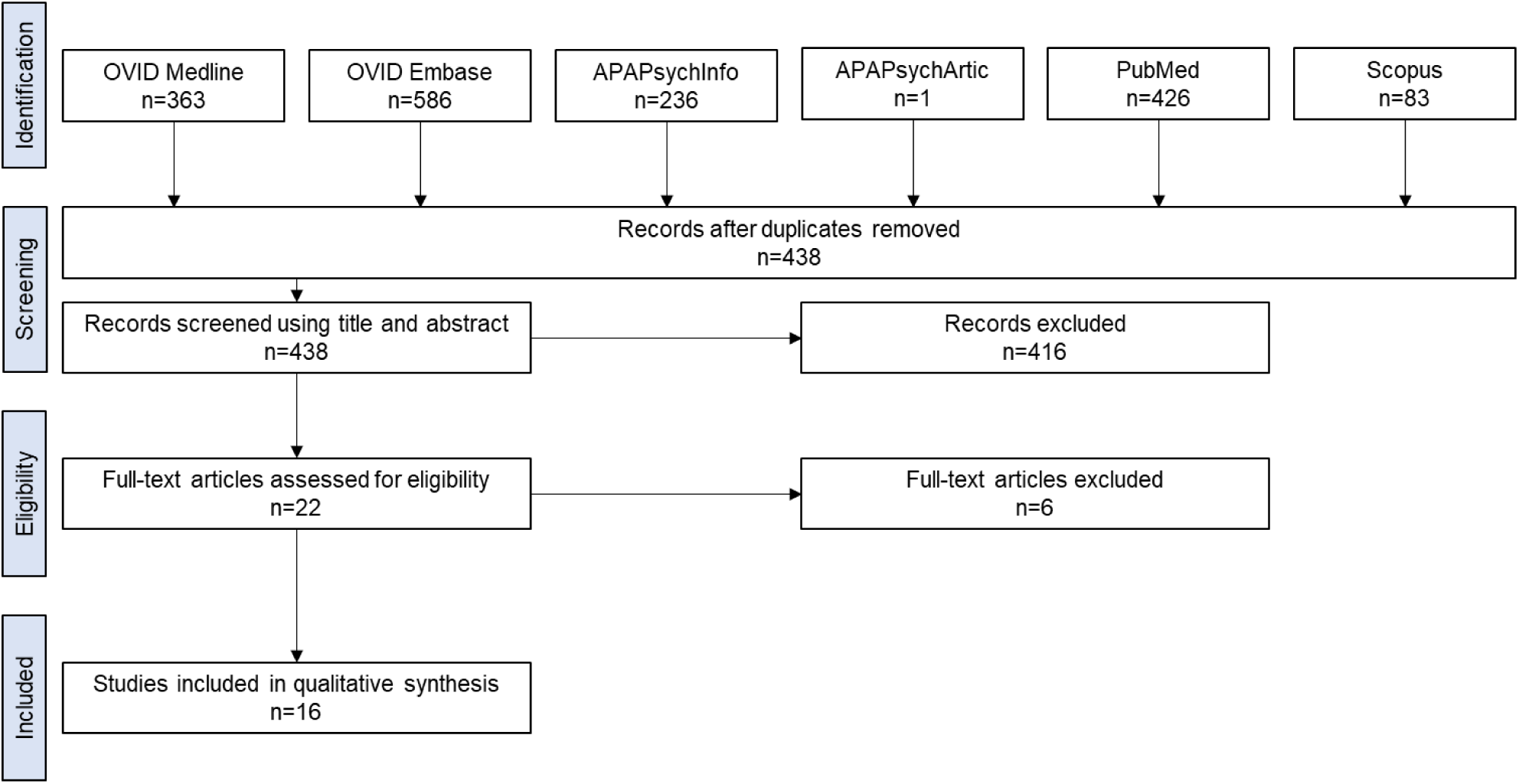
PRISMA diagram outlining search strategy results and article screening process.

For example, a study by Chen et al evaluated differences in coupling between regions in the DMN, ECN, and SN among non-OST HD with short-term (7-15 days) abstinence, long-term abstinence (6-12 months), and HC (Chen et al., 2021). HC showed significantly stronger FC between a core region of the SN, the dorsal Anterior Cingulate Cortex (dACC), to regions in both the ECN (dorsolateral Prefrontal Cortex (dlPFC)) and DMN (precuneus and Posterior Parietal Cortex (PPC)) compared with HD with long-term abstinence. HD with long-term abstinence showed significantly stronger ECN-DMN FC as compared with HD with short-term abstinence (Chen et al., 2021). A later study by the same group replicated this result in participants on short- and long-term OST-maintained abstinence. They showed long-term abstinent and HC participants had stronger FC between the dorsal Anterior Cingulate Cortex (dACC) – dlPFC and dACC – PPC as compared with participants with short-term abstinence (Chen et al., 2022). In both studies by Chen et al, there was a significant, positive relationship between the duration of abstinence and FC strength between regions in the SN - ECN. Additionally, participants with long-term abstinence had significantly lower craving scores than participants with short-term abstinence (Chen et al., 2021, 2022).

Another study also showed significant reductions in craving scores after long-term abstinence in HD, and concurrently, significantly stronger FC in three edges: the Ventral Tegmental Area (VTA) - Orbitofrontal Cortex (OFC), VTA - ACC, and substantia nigra - OFC (Xu et al., 2021). Three additional studies reported both significant reductions in craving and significantly stronger FC after abstinence (Lu et al., 2022; S. Zhang, Yang, Li, Wang, et al., 2022; S. Zhang, Yang, Li, Wen, et al., 2022). Moreover, two out of three studies found a significant relationship between the abstinence-strengthened FC and corresponding decreases in craving (Lu et al., 2022; S. Zhang, Yang, Li, Wang, et al., 2022).

Differences in FC between HC, HD who did relapse, and HD who did not relapse have been evaluated (Q. Li et al., 2018; Wang et al., 2021). Li and colleagues collected baseline rsfMRI scans from HD and HC, as well as the number of relapses per month over a three-month period. The authors found weaker FC between nodes in the ECN (dlPFC) and DMN (mPFC) in HD was significantly associated with a greater number of relapses. Similarly, Wang et al collected baseline rsfMRI data from HD before they began an OST-maintained, 26-month abstinence. The authors found a significant association between the likelihood of relapse and weaker baseline FC between the Nucleus Accumbens (NAc) and regions within the SN and ECN (e.g., the Inferior Frontal Gyrus (IFG) and dlPFC, respectively) (Wang et al., 2021). One study did not find any differences in FC between HU and HC (Xue et al., 2022).

### 3.3 Differences in pairwise resting state functional connectivity changes as a result of TMS between HC and HD

The effects of applying TMS to the dlPFC in HD have been investigated in two studies (Studies 12-13, Table 2). In both studies, HD had a baseline rsfMRI scan, then received randomised TMS (rTMS) to the dlPFC at 10 Hz every day for one week, after which another rsfMRI scan was collected. HC had a rsfMRI scan, but received no intervention. The HC scan was used as a comparison for the before- and after-stimulation rsfMRI scans in HD. Both studies were a part of a larger clinical trial, but it was not clear if any participants were included in both studies. Both studies employed group ICA to identify regions from the DMN, SN, and ECN used in the FC analyses. The first published study by Jin et al showed TMS strengthened left dlPFC - left parahippocampal gyrus FC (i.e., stronger ECN-DMN connectivity), and weakened FC between the right precentral gyrus and three DMN regions: the PCC, praecuneus, and IPL (Jin et al., 2022). The second study by Jin et al showed TMS weakened FC in two edges: the left IPL - right occipital cortex, and the left IPL - left middle frontal gyrus (which is associated with attentional networks (Menon & D’Esposito, 2022; Talati & Hirsch, 2005) (Jin et al., 2023). Despite the similarity in study methods and the focus on regions in the DMN, SN, or ECN, there was no overlap in the edges showing significant changes as a result of TMS. However, both studies showed TMS strengthened FC between regions in the DMN to areas in sensory cortices.

The relationship between TMS-induced changes in FC and measures of spontaneous and drug cue-induced craving was different in each study. The first study showed the stronger dlPFC - parahippocampal gyrus FC was negatively related with craving. In other words, the TMS-strengthened ECN – DMN FC was associated with decreased craving (Jin et al., 2022). The second study found TMS weakened FC in two DMN-SN edges (the IFG – IPL, and IFG – PCC) related to reductions in craving (Jin et al., 2023).

### 3.4 Differences in pairwise resting state functional connectivity changes as a result of acute heroin exposure between HC and HD

Three studies, all of which were conducted by the same group based in Switzerland, investigated the effects of acute heroin administration on FC in HD (Studies 14-16, Table 2). HD FC before and after the acute administration of heroin was compared with FC in HC, who received a placebo/vehicle instead of heroin. All HD must have received heroin-assisted treatment for at least 6 months, with a stable dose for at least 3 months. Two studies administered heroin hydrochloride and collected data during resting state only, whereas in the third, participants also viewed a series of faces with fearful or neutral expressions after heroin dosing. In all studies HD were administered their prescribed dose of heroin hydrochloride in 5 ml of sterile water through an intravenous catheter over 30 seconds. Then, 20 minutes after injection, HD participants performed the MRI scan.

There were no consistent effects of acute heroin administration on FC in HD, nor consistent differences in FC between HD and HC. For example, one of the studies showed that in HD, heroin administration strengthened FC between the putamen and the basal ganglia/limbic regions (Schmidt, Denier, et al., 2015). Conversely, a study by Denier et al showed heroin weakened FC between the thalamus and several regions, including the praecuneus, lateral occipital cortex, superior parietal lobule, middle temporal gyrus, and medial frontal gyrus (Denier et al., 2015).

The study by Denier et al showed most of the significant differences in FC were not between HD and HC, but rather between pre- and post-heroin administration in HD. HC had stronger FC between the thalamus and four regions: the praecuneus, lateral occipital cortex, middle temporal gyrus, and the precentral gyrus. A different relationship was present in the study by Schmidt et al, which showed HC had stronger FC between the PCC/praecuneus and the basal ganglia/limbic network compared with HD at baseline. This difference was abolished under the influence of heroin (Schmidt, Denier, et al., 2015). Administration of heroin also removed differences in FC between HC and HD during an emotional face processing task (Schmidt, Walter, et al., 2015). During the presentation of faces with fearful expressions, heroin administration attenuated the significantly stronger FC in HD pre-administration of heroin vs HC. The weakened FC was between two edges: the left fusiform gyrus - left amygdala, and the right amygdala - right OFC.

## 4. Discussion

An improved mechanistic understanding of how neural processes shape opioid addiction, and how they can be attenuated, informs the development of therapies for opioid addiction. This review intended to develop a better understanding of FC as a neural correlate underpinning opioid addiction. We assessed whether abstinence or interventions affects HD FC, and if changes in FC relate to craving or likelihood of relapse.

We found that HD show differences in FC from HC at baseline. These differences typically consist of weaker FC between regions in the ECN, SN, and DMN (Fig 3). Each of these networks are responsible for a key aspect of healthy brain function. Previous work has shown opioid addiction disrupts DMN, ECN, and SN network function and the processes they mediate. For example, HD have different activity patterns than HC in DMN regions (Pandria et al., 2018; Y. Zhang et al., 2011). DMN regions include the following: mPFC, angular gyrus, superior frontal gyrus, lateral temporal lobes and cortex, temporal junction and pole, the hippocampus, parahippocampal cortex, retrosplenial cortex, PCC, praecuneus, and IPL (Spreng & Andrews-Hanna, 2015). A review of HD vs HC resting state activity related differences in DMN regions to deficits in self-referential processing, emotional regulation, and autobiographical memory (Pandria et al., 2018). HD also exhibit cognitive deficits and atypical function in core ECN regions such as the dlPFC, pre-supplementary motor area and PPC (Lee et al., 2005; K. Shen et al., 2020; Tanabe et al., 2019). The SN includes the right IFG, anterior insula, and the dACC (Menon & Uddin, 2010; Peters et al., 2016). The SN is responsible for the detection of salient stimuli and recruitment of either the DMN or ECN, depending on whether internally-relevant or externally-focused processes is most appropriate, respectively (Menon, 2011). Impairments in the SN’s ability to coordinate DMN and ECN function is associated with deficits reported in HD (Chen et al., 2022; Jin et al., 2022; Q. Li et al., 2018, p. 201). For example, deficits in the SN’s ability to mediate a shift between DMN-controlled internal processes and ECN-mediated, executive control processes impairs goal-directed behaviour (Uddin et al., 2019). Moreover, the rumination and impaired self-awareness in HD are associated with dysfunctional interactions between the DMN, ECN, and SN (Killingsworth & Gilbert, 2010; R. Zhang & Volkow, 2019).

**Figure 3:**
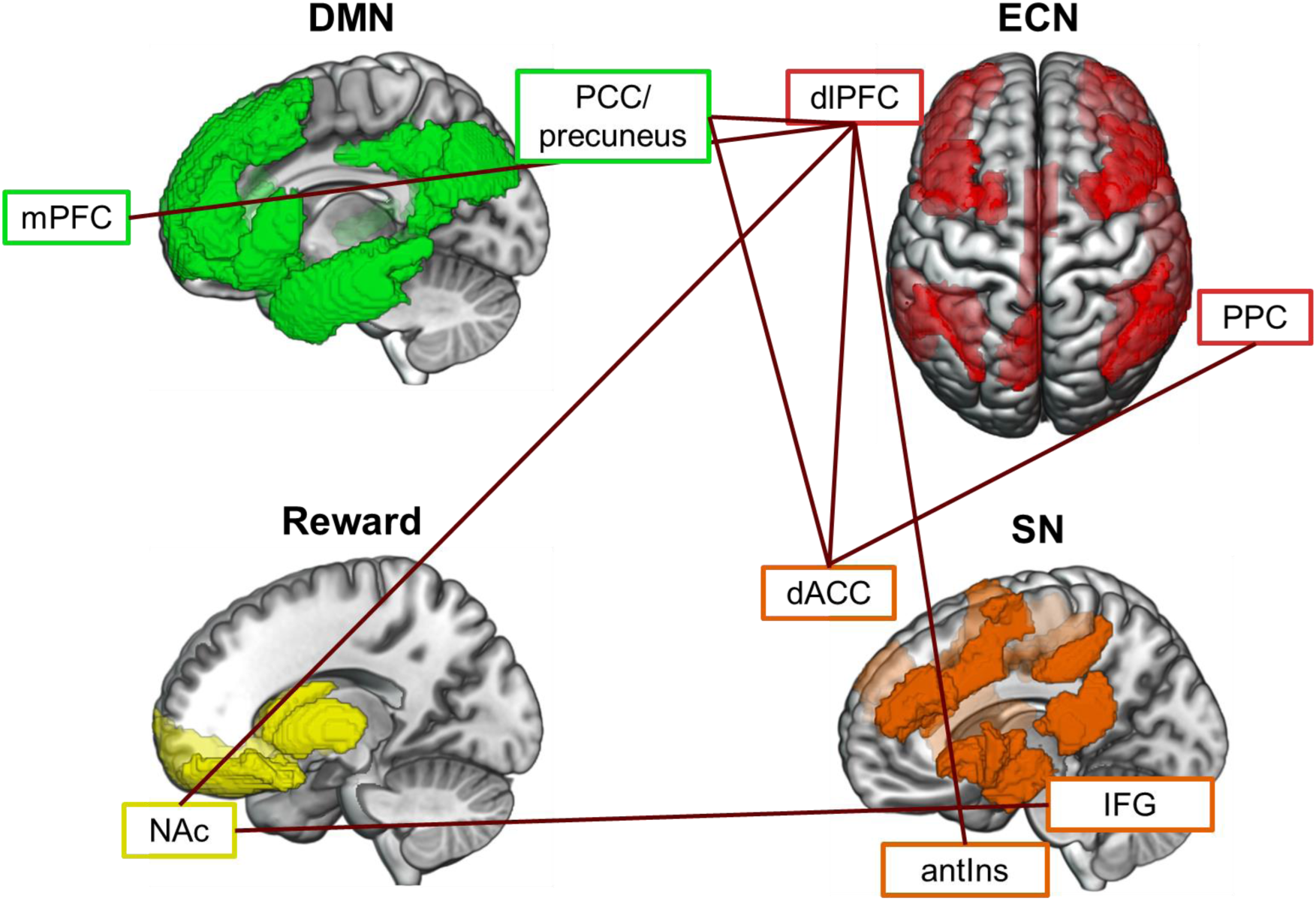
Edges where abstinence significantly strengthened FC in the direction towards HC. Edges (i.e., connections between two regions\) were selected that had replicable relationships across at least two studies and are grouped according to the functional network in which they were categorised. Abbreviations are as follows: mPFC = medial Prefrontal Cortex; PCC = Posterior Cingulate Cortex; dlPFC = dorsolateral Prefrontal Cortex; PPC = Posterior Parietal Cortex; IFG = inferior frontal gyrus; antIns = anterior insula; dACC = dorsal Anterior Cingulate Cortex; NAc = Nucleus Accumbens.

While most HD participants showed weaker baseline FC between the DMN, ECN, and SN than HC, studies consistently reported that abstinence strengthened FC among regions in the ECN, SN, and DMN in the direction towards HC (Fig 4). Additionally, abstinence was shown to relate to reductions in craving and strengthened FC among reward areas (Lu et al., 2022; Xu et al., 2021), and from reward areas to SN (Wang et al., 2021; Xu et al., 2021) and ECN (Wang et al., 2021) (Fig 4). While one study showed abstinence strengthened (rather than weakened) FC among ECN, SN, and DMN regions (S. Zhang, Yang, Li, Wang, et al., 2022), the change in FC was otherwise consistent in that abstinence shifted FC towards the stronger FC observed in HC vs HU. This accords with work showing that HD have weaker FC as compared with HC between key regions in the ECN (e.g., dlPFC) and DMN (e.g., IPL) (Yuan et al., 2010), and that abstinence strengthens FC among DMN, ECN, or SN regions (Yang et al., 2022).

**Figure 4:**
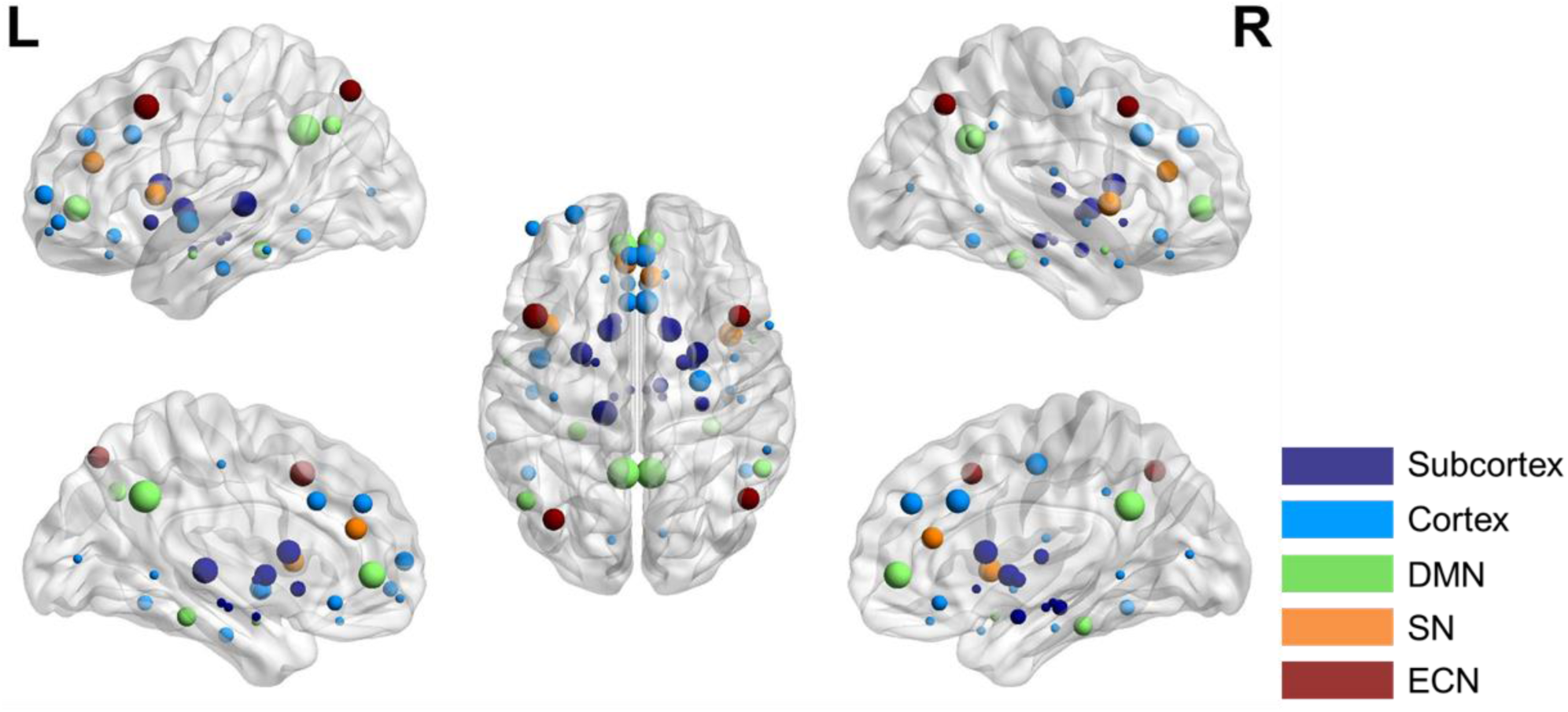
Glass brains showing ROIs with different FC between HC and HD as a result of an abstinence, TMS, or acute opioid exposure. The size of the region scales with the number of times it was mentioned across studies. The colour of each region indicates whether a study grouped it within the DMN, SN, or ECN. All other regions were grouped into subcortical or cortical groups. This figure was made using BrainNet (Xia et al., 2013).

The consistency of the effect of abstinence on FC in HD is notable since studies utilising both OST- and non-OST-supported abstinence were included in this work. OST-supported abstinence is different from opioid-free abstinence. OST has been proposed to maintain the hypodopaminergic state/trait in HD (Blum & Baron, 2019) which contributes to aberrant, reward-related neural processes associated with addiction. Given that both OST- and non-OST-supported abstinence tended to strengthen HD FC towards that seen in HC, stability in opioid use may relate to a “normalisation” of FC patterns. In turn, the stability-associated changes in FC may relate to reductions in craving and relapses.

**Figure 5:**
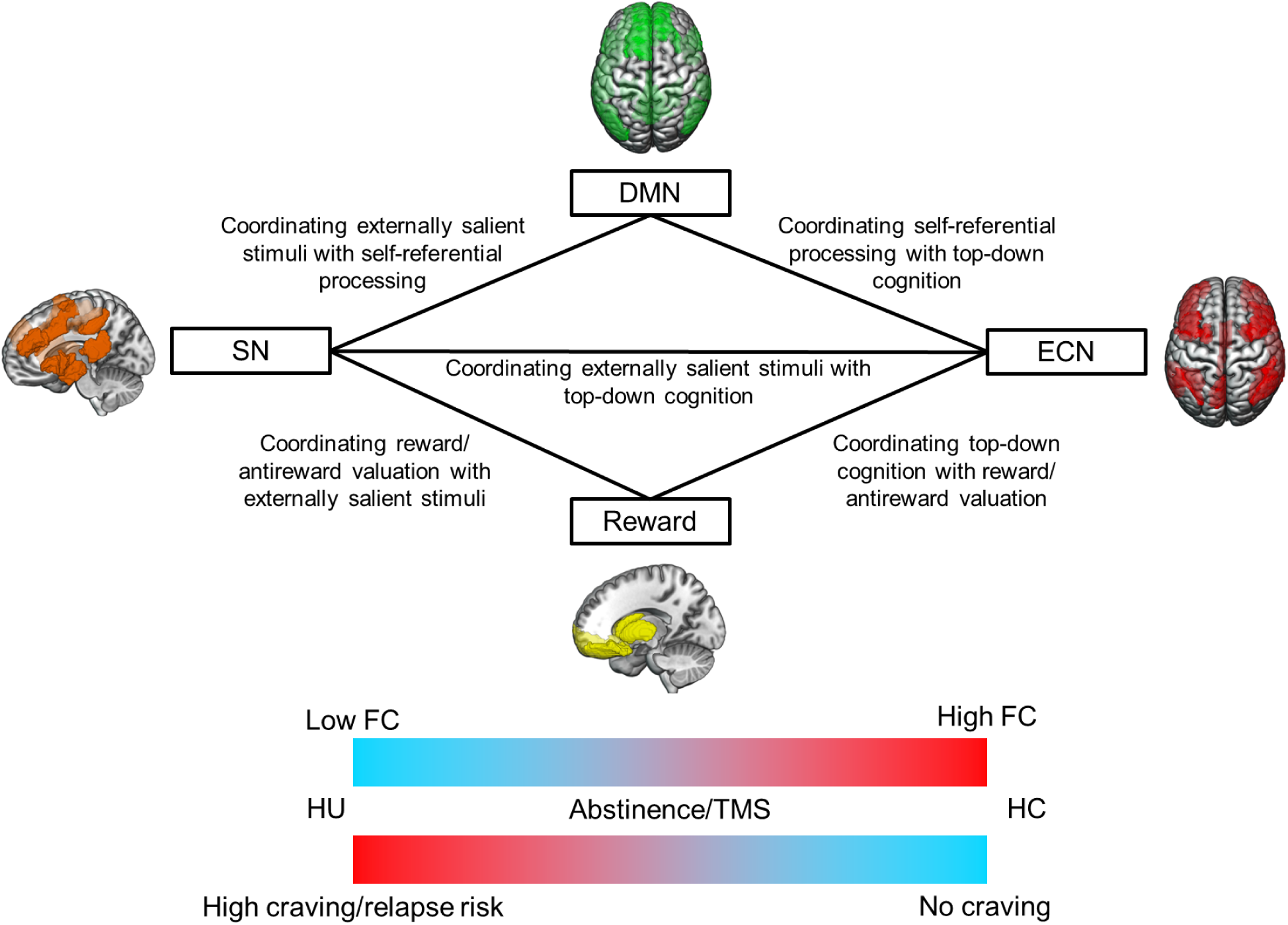
Schematic of how weakened FC introduces deficits in processes mediated by between-network connectivity, and its relationship to the risk of craving and relapse.

Similar to the effects of abstinence on FC and craving/relapse, studies showed that TMS strengthened FC of ECN, SN, and DMN regions in HD towards HC, which related to decreased craving and relapse (Fig 4). While the evidence supporting a clinically relevant relationship between craving and relapse is unclear (Hasin et al., 2013; Tiffany & Wray, 2012), the effects of abstinence on craving and relapse reviewed here (Q. Li et al., 2018; Wang et al., 2021) suggest that reduced craving is associated with reduced likelihood of relapse. This aligns with work showing decreased craving reduces the risk of relapse in HD (Saraiya et al., 2021; Tsui et al., 2014), and that stronger FC among the ECN, SN, and DMN predicted lower risk of relapse in people with opioid, cocaine, methamphetamine, nicotine, and alcohol addiction (Moeller & Paulus, 2018).

The effects of TMS on craving reviewed here also accord with other investigations of the effects of TMS on craving in HD (X. Li et al., 2021; X. Liu et al., 2020; Y. Shen et al., 2016). However, a study investigating whether TMS of the dlPFC reduces craving in HD did not show a reduction in craving (Tsai et al., 2021). While a review posited this discrepancy was due to the inclusion (Tsai et al., 2021) or exclusion (X. Li et al., 2021; X. Liu et al., 2020; Y. Shen et al., 2016) of participants on OST (Mehta et al., 2024), the two studies reviewed in this work showing TMS-induced reductions in craving included participants on OST (Jin et al., 2022, 2023). Moreover, another review of the effects of neuromodulation on craving in opioid addiction included three TMS studies (Ward et al., 2020), two of which had not been included in the review by Mehta et al. While neither study reported whether their study in- or excluded participants on OST, and one study did not target the dlPFC (Y. Shen et al., 2017), both studies showed decreases in craving following TMS (Sahlem et al., 2017; Y. Shen et al., 2017). While replicability across neuromodulation studies is often low due to high experimenter and interindividual variability (Guerra et al., 2020; L. M. Li et al., 2015), the post-TMS reduction in craving in HD shown in seven out of eight TMS studies encourages further exploration of TMS as a method to reduce craving and relapse.

Networks that underpin stress, reward, and emotional circuitry are also affected by opioid addiction. These networks, referred to as the limbic and ventromedial networks, include regions such as the NAc, medial OFC, mPFC, and midbrain structures such as the caudate, putamen, and amygdala (Dunlop et al., 2017; Everitt & Robbins, 2016; Koob & Volkow, 2010). Together, these regions encode positive or negatively valanced events/stimuli, incentive drive, and stress responses. For a comprehensive description of the effects of addiction on reward/emotional circuitry we refer the reader to the following work: (Everitt & Robbins, 2016; Gold et al., 2020; Koob & Volkow, 2010). Reward network regions may also be associated with the DMN, SN, or ECN, as the role of a brain region within a network is context-dependent (Cole et al., 2013; Matthews & Hampshire, 2016; Medaglia et al., 2015; Soreq et al., 2019). For example, the mPFC is considered a core region of both the DMN and reward networks. During rest, the mPFC’s function will be associated with other DMN regions, whereas during a reward-related task, it will be associated with regions in reward networks (Spreng & Andrews-Hanna, 2015; Tzschentke, 2000; van Holstein & Floresco, 2020). Yet, the mPFC was only considered as a DMN region in the studies included in the review. It would have been informative to see analyses investigating whether the mPFC had stronger FC to other regions in the DMN or reward-related networks. Such an analysis would enable greater sensitivity in characterising whether the role of the changes in mPFC FC are associated with either self-referential (e.g., DMN-related) or reward processes disrupted by opioid addiction.

It was notable that all studies investigating the effects of abstinence or acute opioid administration on FC were conducted in either China or Switzerland, respectively. Studies conducted solely in one country may not be representative of the broader population (Ge et al., 2023; Han & Northoff, 2008; Kitayama & Salvador, 2017). For example, abstinence programs that differ across countries may introduce variability in the neural markers of abstinence. Some countries (e.g., China) utilise compulsory isolated treatment, whereas other countries (e.g., US, UK) typically do not (National Institute for Health Care and Excellence (NICE), 2007; Tibke, 2017). Similarly, the administration of heroin was in line with Switzerland’s therapeutic strategy for HD, which utilises pharmaceutical heroin diacetylmorphine (DAM). While these studies align with clinical practice in Switzerland, it is important to conduct more international research on the neural correlates of addiction and their response to treatment. Such work would substantiate whether strengthened FC is a suitable biomarker of responses to treatment for opioid addiction.

The methods employed in a study impact results and interpretation (Box 1, Box 2). While most studies evaluating the effect of abstinence on FC in HC and HD defined their regions using standard functional or anatomical parcellations, three studies (Wang et al., 2021; Wei et al., 2021; Xue et al., 2022) chose regions based on the presence of statistically significant differences in: regional homogeneity between HC, short, and long-term abstaining people with OUD (Xue et al., 2022), voxelwise differences in Degree Centrality or BOLD activity between HC and OUD participants (Wang et al., 2021), and voxelwise activity when viewing heroin vs neutral cues (Wei et al., 2021). The purpose of selecting regions of interest based on voxelwise measures of significance is to narrow the scope of analysis within the vast quantity of data generated by fMRI. However, this practice is referred to as “double-dipping”, and is a form of circular analysis (Kriegeskorte et al., 2009). Circular analyses often occur in large, noisy datasets (e.g., fMRI data), where a selection of the data that is significant in one analysis is used in subsequent statistical tests. The noise contained in the first analysis is amplified in the second, increasing the likelihood of a type I error, i.e., a false positive. To avoid this issue, it is recommended to use independent study populations, or better yet, meta-analysis, to identify regions of interest (Box 1). ICA-derived regions are another robust way to focus analyses on regions shown to be reliably engaged in the condition (e.g., task or resting state) of interest without assuming statistical differences between groups (Duff et al., 2012). To summarise, the parcellation method by which regions of interest are selected is an important part of FC analyses and may influence the quality of results. Readers are advised to consult Kriegeskorte’s guidelines on selecting regions of interest to reduce the impact of noise and bias in their results (Kriegeskorte et al., 2009).

Another methodological comment concerns the disparity in clinical factors (e.g., sex, socioeconomic status, neurodiversity) in study populations. For example, nearly half of the studies in this review only included men. There is some literature showing FC differences between HC males and females (Weis et al., 2020), and there are some sex-based differences in the functional correlates of addiction (Nicolas et al., 2022; Smith et al., 2023). Similar effects of neurodiversity and socioeconomic status on FC have been reported (Horien et al., 2023; Sripada et al., 2021; Yaple & Yu, 2020). Therefore, future work should endeavour to include representation of male and female participants, as excluding the consideration of clinically relevant factors from research negatively impacts the clinical relevance of findings (Johnson et al., 2014).

### 4.1 Conclusions

There is a pressing need to characterise the mechanisms of addiction and develop therapies that attenuate them. Recent studies delivering noninvasive brain stimulation guided by activity-based analyses provides an example of a successful, research-informed therapy. These studies showed administering noninvasive brain stimulation to personalised anatomical targets during an individualised cue-reactivity paradigm substantially decreased craving and substance use in HD (Mahoney, Haut, et al., 2023; Mahoney, Thompson-Lake, et al., 2023). FC can extend the insights garnered by activity-based analysis and identify a clearer link between targets for therapeutic interventions and the quantification of the therapies’ success. This work suggests that either abstinence or TMS changes FC in HD in the direction of HC, which in turn, relates to reduced craving and risk of relapse. Improving the methodological practices and widening the distribution of research conducted in this field will not only strengthen the robustness of these findings, but also facilitate the application of FC analyses to translational research and clinical practice.

## Appendix

### Search strategies

#### PubMed

(“functional connectivity”[Title/Abstract] OR “functional magnetic resonance imaging”[Title/Abstract] OR “fMRI”[Title/Abstract] OR “functional imaging”[Title/Abstract] OR “functional-magnetic”[Title/Abstract] OR “connectivity”[Title/Abstract] OR “connectome”[Title/Abstract]) AND (”oud”[Title/Abstract] OR “use disorder”[Title/Abstract] OR “addiction”[Title/Abstract] OR “addict*”[Title/Abstract] OR “dependen*”[Title/Abstract] OR “behavior, addictive”[MeSH Terms] OR “addiction medicine”[MeSH Terms] OR “substance related disorders”[MeSH Terms] OR “substance abuse, intravenous”[MeSH Terms] OR “substance abuse, oral”[MeSH Terms] OR “opiate substitution treatment”[MeSH Terms]) AND (”opiu*”[Title/Abstract] OR “heroin”[Title/Abstract] OR ((”illicit drug”[Title/Abstract] OR “illicit drugs”[MeSH Terms]) AND “opio*”[Title/Abstract]) OR “opium”[MeSH Terms] OR “heroin”[MeSH Terms] OR “heroin dependence”[MeSH Terms] OR “opium dependence”[MeSH Terms])

#### OVID: MEDLINE, EMBASE, PsycINFO, PsycArticlesFullText

1. opio*.ti. or opio*.ab. 304848
2. illicit.ti. or illicit.ab. 59219
3. opiu*.ti. or opiu*.ab. or heroin.ti. or heroin.ab. 54390
4. (opium or opium dependence or opioid-related disorders or heroin or heroin dependence).mh. 10078
5. opium/ or opium dependence/ or heroin/ or heroin dependence/ 172040
6. addictive behaviour/ or exp addiction medicine/ or substance abuse/ or substance abuse, intravenous/ or substance abuse, oral/ 284523
7. functional connectivity.ab. or functional connectivity.ti. or fMRI.ab. or fMRI.ti. or functional magnetic resonance imaging.ab. or functional magnetic resonance imaging.ti. or functional-magnetic.ab. or functional-magnetic.ti. or functional imaging.ab. or functional imaging.ti. or connectivity.ab. or connectivity.ti. or connectome.ab. or connectome.ti. 380338
8. oud.ab. or oud.ti. or “use disorder”.ab. or “use disorder”.ti. or addic*.ab. or addic*.ti. or dependen*.ab. or dependen*.ti. or (Behavior, Addictive or Addiction Medicine or Substance-Related Disorders).mh. 4707938
9. (heroin or heroin dependence).mh. 4403
10. 1 and 2 8595
11. 3 or 4 or 5 or 10 201828
12. 6 or 8 4886002
13. 7 and 11 and 12 666
14. remove duplicates from 13 434

#### SCOPUS

(((TITLE(opiu*) OR ABS(opiu*) OR TITLE(heroin) OR ABS(heroin) OR ((TITLE(opio*) OR ABS(opio*)) AND (TITLE(illicit) OR ABS(illicit))) AND ((TITLE(functional AND connectivity) OR ABS(functional AND connectivity) OR TITLE(fmri) OR ABS(fmri) OR TITLE(functional AND magnetic AND resonance AND imaging) OR ABS(functional AND magnetic AND resonance AND imaging) OR TITLE(functional-magnetic) OR ABS(functional-magnetic) OR TITLE(functional AND imaging) OR ABS(functional AND imaging) OR TITLE(connectivity) OR ABS(connectivity) OR TITLE(connectome) OR ABS(connectome))) AND ((TITLE(oud) OR ABS(oud) OR TITLE(”use disorder”) OR ABS(”use disorder”) AND TITLE(addic*) OR ABS(addic*) OR TITLE(dependen*) OR ABS(dependen*)))

